# *miR17~92* is essential for the survival of hematopoietic stem and progenitor cells by restraining pro-apoptotic BIM

**DOI:** 10.1101/342071

**Authors:** Kerstin Brinkmann, Craig Hyland, Carolyn A de Graaf, Andreas Strasser, Warren S Alexander, Marco J Herold

## Abstract

The micro RNA cluster *miR17~92*, also known as oncomiR-1, impacts diverse cellular processes, such as cell survival and proliferation. Constitutive loss of *miR17~92* in mice causes severe defects in skeletal development, organ development and hematopoiesis, resulting in early post-natal lethality. The critical functions of *miR17~92* in a fully developed animal have not yet been explored. Here we show that deletion of *miR17~92* in adult mice 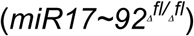 had no impact on their lifespan or general well-being. However, detailed analysis of the hematopoietic system in 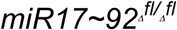 mice, revealed a dramatic reduction in all mature hematopoietic lineages, which was due to the loss of early hematopoietic stem/progenitor cells (HSPCs). Strikingly, the concomitant loss of the pro-apoptotic BH3-only protein BIM rescued the loss of the HSPCs and all of their differentiated progeny that was caused by the deletion of *miR17~92*. These findings demonstrate that *miR17~92* is critical for the survival of HSPCs by restraining the activity of the pro-apoptotic BH3-only protein BIM.

## Introduction

The *miR17~92* cluster comprises six micro RNAs, including *miR17*, *miR18a*, *miR19a*, *miR20a*, *miR19b1* and *miR92-1*, which belong to 4 different seed families ^1^. The *miR17~92* cluster is also known as *oncomiR-1*, due to its well-described function as an oncogene ^1^. Specifically, *miR17~92* is over-expressed in certain B cell lymphomas, including diffuse large B cell lymphoma, T cell lymphomas, acute myeloid leukemia (AML), chronic lymphocytic leukemia (CLL) and a range of solid tumors, including retinoblastoma, neuroblastoma, osteosarcoma as well as cancers of the colon, pancreas, breast, ovaries, lung, kidney and liver (reviewed in ^2^). Notably, *miR17~92* over-expression is linked to poor prognosis in diverse cancers (reviewed in ^2^). Accordingly, transgenic studies in mice revealed a role for *miR17~92* in MYC-driven lymphomagenesis (*Eμ-Myc* mouse model) and in prostate cancer development ^3-6^. MYC increases transcription of the *miR-17-92* cluster ^7^, and this is thought to promote tumorigenesis through repression of several genes involved in cell cycle regulation (e.g. *Pten*, *E2F1-3*)^8-10^, angiogenesis (e.g. *Tsp1*)^11^, TGFb receptor signaling (e.g. *TGFBRII*, *Smad2*, *Smad4*)^12^ and apoptosis (e.g. *Bim)* ^13^.

Studies with mice constitutively deficient for *miR17~92* revealed that this micro RNA cluster is essential for the normal development of several organs, including the heart, lung and skeleton as well as for the normal production of B and T lymphoid cells ^10,13-16^. The *miR17~92^-/-^* mice die soon after birth with severe lung and ventricular septal defects ^13^. Moreover, at birth the *miR17~92^-/-^* mice were significantly smaller in size compared to their wild-type littermates and presented with severe skeletal abnormalities, similar to the defects observed in the human Feingold syndrome ^17^. The inducible (conditional) deletion of the *miR17~92* cluster using tissue specific CRE transgenes showed that *miR17~92* plays a critical role in the development of several stem and progenitor cell populations, including those for osteoclasts, nephrons, neuronal cells and endothelial cells ^18-21^.

Adult wild-type mice that had been lethally irradiated and then had their hematopoietic compartment reconstituted with stem/progenitor cells from the fetal liver of *mir17~92^-/-^* embryos (E14.5) presented with abnormally low numbers of circulating B cells, splenic B cells and pre-B cells in the bone marrow ^13^. Consistent with a critical role for the *miR17-92* cluster in early B lymphopoiesis, a substantial reduction in pro-B/pre-B cells was also observed in the fetal liver of E18.5 *miR17~92^-/-^* embryos ^13,17^. Conversely, overexpression of the *miR-17-92* cluster selectively in lymphocytes (*miR17~92^tg/tg^;hCD2-iCre* mice) caused lymphoproliferative and autoimmune disease ^10^, which is reminiscent of the phenotypes seen in mice deficient for pro-apoptotic BIM ^22^ or transgenic mice over-expressing pro-survival BCL-2 in the B cell lineage ^23^. Hematopoietic cell specific loss of *miR-17*~*92* (*miR17~92^fl/fl^;vav-iCre* mice) resulted in profound defects in T cell development both at the level of T cell progenitors in the thymus and at later stages of differentiation ^24^. Conversely, retroviral-mediated *miR17~92* over-expression promoted expansion of multi-potent hematopoietic progenitors and significantly increased the colony forming capacity of mouse bone marrow progenitor cells *in vitro* ^25^.

Collectively, these findings are consistent with the notion that the defects in the hematopoietic system caused by the loss of *miR17~92* might be a consequence of a loss of the HSPC population rather than isolated effects on different hematopoietic cell populations ^26^. To date, no studies have investigated the impact of the inducible organism-wide deletion of *miR17~92* in a fully developed animal. Furthermore, the target(s), derepression of which is critical for the cellular defects caused by the loss of the *miR-17-92* cluster remains to be defined, with currently no genetic proof of *in vivo* relevance of any functional interaction. This is surprising given that the *miR17~92* cluster is considered a promising target for the therapy of cancer and certain other diseases. Here we report the impact of organism-wide induced deletion of the *miR17~92* cluster in adult mice. This study reveals that such loss of *miR17~92* does not impact the overall well-being of mice but causes a severe depletion of various hematopoietic stem and progenitor cell populations. Remarkably, all of these defects can be fully prevented by the concomitant loss of the pro-apoptotic BH3-only protein BIM.

## Results

### Inducible deletion of *miR17~92* in adult mice causes a substantial loss of diverse hematopoietic cell types

We wanted to determine the critical function of *miR17~*92 in adult mice. To this end we generated *miR17~92^fl/fl^Rosa-CreERT2^Ki/wt^* mice^27^ and treated them alongside control *Rosa26CreERT2^Ki/wt^*, *miR17~92^fl/fl^* as well as C57BL/6 wild-type (wt) animals (males and females, aged 8-12 weeks) for 3 days with tamoxifen and observed them for another 180 days.

The CRE-induced deletion of the *miR17~92* cluster in adult mice led to no obvious abnormalities in behavior, breathing, food intake, weight loss or severe anemia, which served as general markers for normal function of the cardiovascular, digestive and blood systems, respectively. This is in striking contrast to the constitutive loss of *miR17~92* from conception, which causes the death of mice soon after birth ^13^.

However, as predicted from previous reports ^10,13,17,24,25^, a significant loss of various hematopoietic cell populations was observed upon deletion of *miR17~92* in the adult mice. This was evident from a loss of lymphocytes, red blood cells, platelets as well as myeloid cell populations, including neutrophils, eosinophils and monocytes (Figure 1a and Supplementary Figure 1a). The reduction of mature red blood cells manifested in a mild anemia as shown by a decrease in the hematocrit (HCT) and hemoglobin (HGB) content (Supplementary Figure S1b). Flow cytometric analysis revealed abnormally low splenic cellularity due to reduced numbers of B cells, T cells, monocytes/macrophages and granulocytes. A reduction in the B cell populations (including pro-B/pre-B cells, immature, transitional and mature B cells) was also observed in the bone marrow of tamoxifen treated *miR17~92^fl/fl^Rosa-CreERT2^Ki/wt^* mice (Supplementary Figure S1c). However, no changes in the architecture of the bone morrow and spleen, were evident in these animals (Supplementary Figure S1d). Interestingly, we observed more severe defects one month after tamoxifen-induced deletion of *miR17~92* compared to animals analyzed 6 months post treatment (Figure 1a and Supplementary Figure S1a, b, c). This raised the possibility that the milder effects observed at the later time points are the result of selective outgrowth and accumulation of cells that had escaped *miR17~92* deletion in the tamoxifen treated *miR17~92^fl/fl^Rosa-CreERT2^Ki/wt^* mice. Indeed, the deletion of *miR17~92* in the hematopoietic tissues (bone marrow, spleen) was almost complete on day 7, but less than 50% in most mice 6 months post tamoxifen administration (Figure 1b and Supplementary Figure S1e). In line with previous reports ^28^, the induction of CRE recombinase by tamoxifen treatment caused measurable toxicity to the hematopoietic cells at early time points but this was overcome within ~35 days post treatment (Figure 1a and Supplementary Figure S1a, b, c). Notably, at this time point the efficacy of *miR17~92* deletion was already less than 50% in the bone marrow as well as spleen of *miR17~92^fl/fl^;RosaCreERT2^Ki/wt^* mice (Figure 1b), suggesting a competitive disadvantage of *miR17-92*-deleted cells.

**Figure 1:**
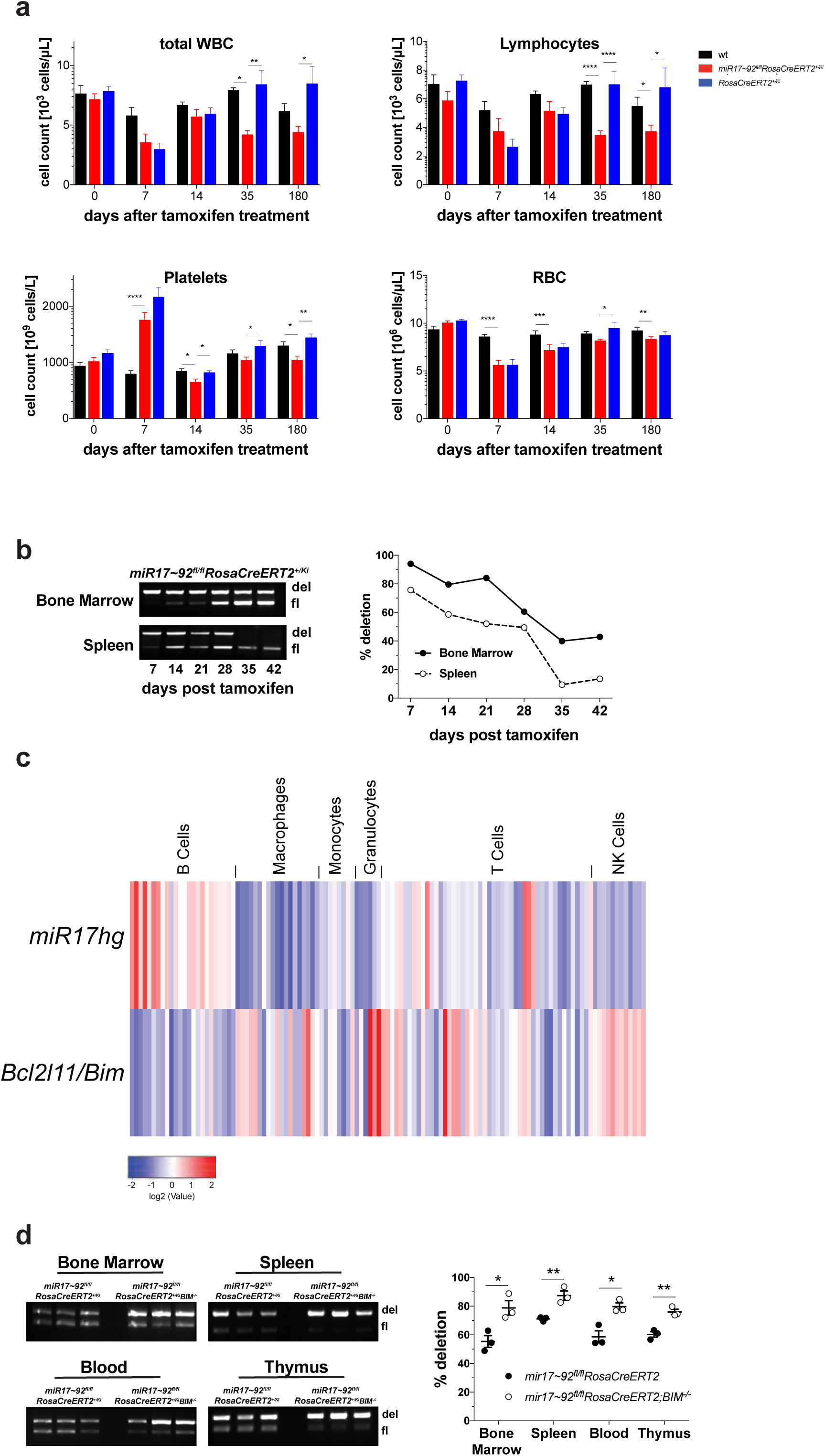
Induced deletion of *miR17~92* causes a substantial loss of diverse hematopoietic cell types in adult mice. **(a)** Mice of the indicated genotypes were treated with tamoxifen to cause activation of the latent CreERT2 recombinase and resultant recombination of the floxed *miR17~92* alleles. Mandible bleeds were taken for hemograms at the indicated time points to determine the numbers of white blood cells (WBC), lymphocytes, platelets and red blood cells (RBC). **(b)** Genotyping (detecting *miR17~92* wt, floxed, del bands) of bone marrow (top left) and spleen cells (bottom left) from *miR17~92^fl/fl^;RosaCreERT2^Ki/wt^* mice at the indicated time points after CreERT2 activation by treatment with tamoxifen (left panel). Deletion efficacy was calculated by densitometry of the PCR products of the deleted *vs* the non-deleted alleles (right panel). **(c)** Correlation analysis of *mir17hg (miR17~92*) and *BCL2L11 (Bim)* RNA expression in the indicated cell types was determined by the examination of microarray data provided by immgen.org ^29^. **(d)** Genotyping (detecting *miR17~92* wt, floxed, del bands) of cells from the bone marrow, spleen, blood and thymus of *miR17~92^fl/fl^;RosaCreERT2^Ki/wt^* (n=3) and *miR17~92^fl/fl^; RosaCreERT2^Ki/wt^;Bim^-/-^* mice (n=3) 3 months after CreERT2 activation by treatment with tamoxifen (left panel). Deletion efficiency was calculated by densitometry of the PCR products of the deleted *vs* the non-deleted alleles (right panel).

### The loss of *miR17~92*-deleted hematopoietic cells can be prevented by the concomitant deletion of BIM

Since BIM expression has previously been shown to increase in pro-B cells upon deletion of *miR17~92* ^13^, we tested whether the expression levels of *Bim* mRNA and the *miR17~92* cluster also correlated inversely in other hematopoietic cell types. Strikingly, microarray analysis obtained through the Immgen database (https://www.immgen.org)^29^ revealed a negative correlation of *Bim (BCL2L11)* and the *miR17~92* cluster host gene *(miR17hg)* expression in all immature and mature lymphocytes, several progenitor populations and mature cell types of myeloid origin, including macrophages, monocytes and granulocytes (Figure 1c).

This prompted us to test whether the deletion of BIM in the *miR17~92^fl/fl^RosaCreERT2^Ki/wt^* mice could restore normal numbers of *miR17~92* deleted hematopoietic cells, by generating *miR17~92^fl/fl^;RosaCreERT2^Ki/wt^BIM^-/-^ mice*. Firstly, we analyzed the deletion efficiency of the *miR17~92* cluster in hematopoietic organs of these mice. While the efficacy of *miR17~92* deletion in hematopoietic cells was significantly less than 50% at 3 months post tamoxifen treatment in the *miR17~92^fl/fl^;RosaCreERT2^Ki/wt^* mice, almost complete deletion was observed in cells from bone marrow, spleen, blood or thymus of the *miR17~92^fl/fl^;RosaCreERT2^Ki/wt^;Bim^-/-^* mice (Figure 1d). These data suggest that loss of BIM may be sufficient to rescue the survival defects of hematopoietic cells that had lost *miR17~92*.

### Loss of pro-apoptotic BIM prevents the reduction in lymphoid, myeloid and erythroid cells that is caused by the induced deletion of *miR17~92*

Our data suggest that BIM may be critical for the loss of the hematopoietic cell populations that occurs upon the induced deletion of the *miR17~92* cluster. To examine the functional interaction of *miR17-92* and BIM-mediated apoptosis in these cell populations, we performed mixed bone marrow reconstitution assays. Lethally irradiated wild-type (wt; C57BL/6-Ly5.1) mice were reconstituted with a 1:1 mixture of GFP-expressing wt (GFP-C57BL/6-Ly5.2) competitor bone marrow cells and test bone marrow cells of the genotype of interest (e.g. *miR17~92^fl/fl^;RosaCreERT2^Ki/wt^*). After confirming adequate 1:1 hematopoietic reconstitution by competitor and test cells by FACS analysis of blood cells ~10 weeks post-transplantation, recipient mice were treated with tamoxifen to delete the *miR17~92* cluster in the test cells (experimental design depicted in Supplementary Figure S2). This allowed us to perform side-by-side comparison of control cells (*miR17~92*-sufficient; GFP expressing wt cells) and cells that had just lost *miR17~92* (*miR17~92^fl/fl^;RosaCreERT2^Ki/w^*) or had lost *miR17~92* on a BIM deficient background (*miR17~92^fl/fl^;RosaCreERT2^Ki/wt^;Bim^-/-^*). As predicted, 60 days post tamoxifen treatment there were significant reductions in several hematopoietic cell populations (e.g. immature and mature B and T cells, myeloid cells) that had lost the *miR17~92* cluster, while wt as well as *RosaCreERT2^Ki/wt^* cells were not outcompeted by the GFP-expressing wt cells (Figure 2). Similar results were obtained when transplanting *miR17~92^-/-^* HSPCs derived from E14 fetal livers in competition with GFP-expressing HSPCs (Supplementary Figure 3). Remarkably, deletion of *miR17~92* on a BIM-deficient background did not impact their competitiveness and resulted in at least a 50% contribution of 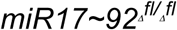;*Bim^-/-^* and GFP-expressing cells to the lymphoid and myeloid cell subsets (Figure 2). In fact, their contributions to the B lymphoid cells in the bone marrow as well as the mature B and T cells in the spleen were significantly greater than 50%, consistent with the accumulation of these cell types seen in BIM-deficient mice ^22^.

**Figure 2.**
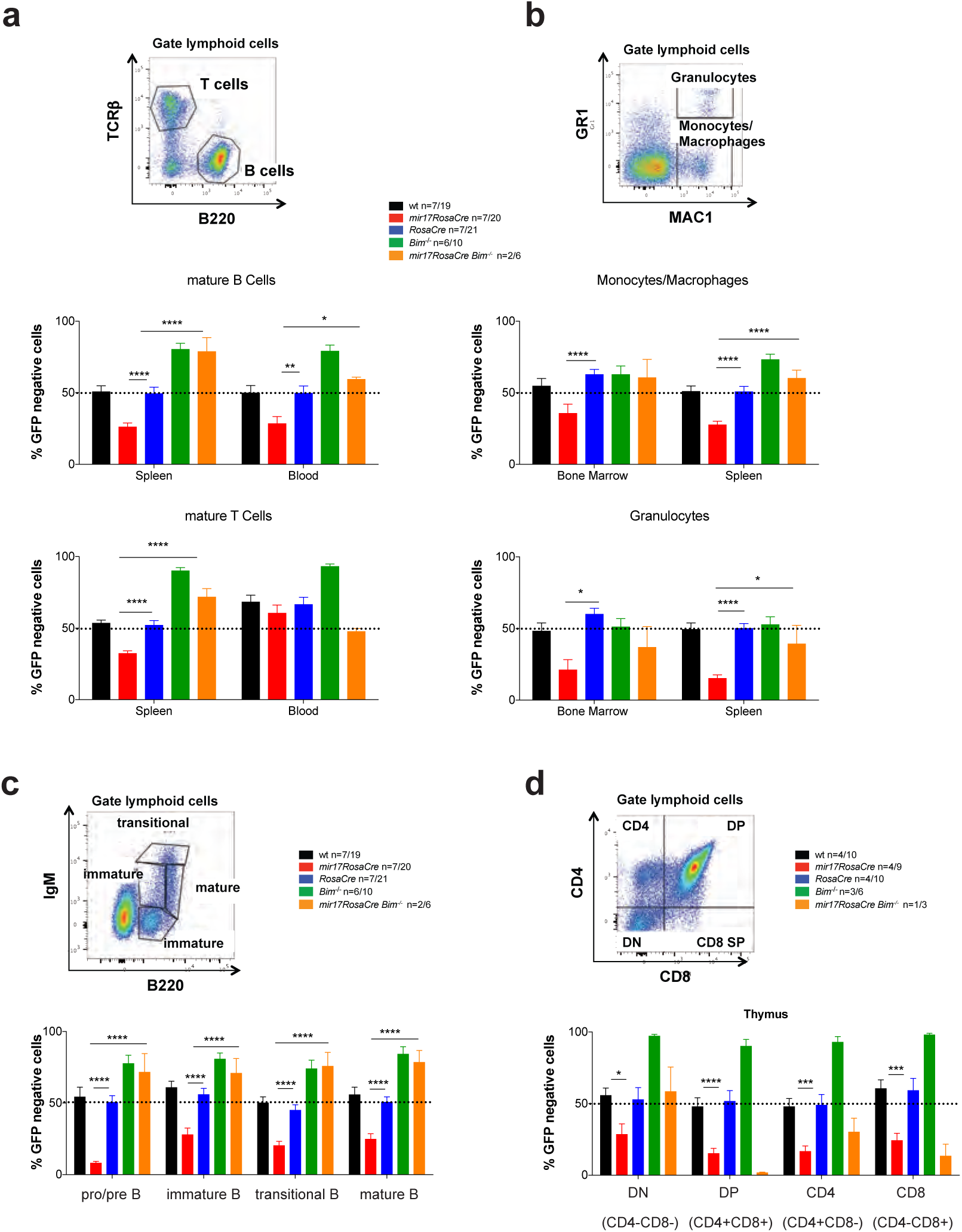
Loss of BIM rescues hematopoietic cells with deleted *miR17~92*. **(a-d)** C57BL/6 wt mice (Ly5.1+) were lethally irradiated and reconstituted with a 1:1 mixture of bone marrow cells from UBC-GFP mice (competitor cells) and bone marrow cells from mice of the indicated genotypes (test cells). Reconstituted mice were treated with tamoxifen (3 doses oral gavage, 60 mg/kg/day) 8 weeks post-transplantation to activate the latent CreERT2 recombinase. After an additional 8-10 weeks, the immature as well as mature lymphoid and myeloid cell populations indicated were analyzed by flow cytometry. **(a)** Mature B and T cells in the spleen and peripheral blood were identified by staining with antibodies against B220 and TCRβ. **(b)** Monocytes/macrophages (MAC-1^+^GR-1^low^) and granulocytes (MAC-1^+^GR-1^high^) were identified in the bone marrow and spleen by flow cytometry**. (c)** Immature and mature B cell populations of the bone marrow were stained with antibodies against B220 and IgM and pro-B/pre-B (B220^+^IgM-), immature B (B220^low^IgM^+^, transitional B (B220^+^IgM^high^) and mature B cells (B220^high^IgM^+^) were identified. **(d)** Immature and mature thymic T cell populations were identified by staining with antibodies against CD4 and CD8. DN=double negative CD4-8-: DP=double positive CD4+8+ cells. Representative FACS blots for the gating strategies are provided for wt mice. Percentages of GFP-negative cells (competitor cells) were determined for each cell subset. Data represent mean +/-SEM. *p<0.05, **p<0.01, ***p<0.001, ****p>0.0001 (Student t test comparing *miR17~92^fl/fl^;RosaCreERT2^Ki/^* with *RosaCreERT2^Ki/wt^* and *miR17~92^fl/fl^;RosaCreERT2^Ki/wt^;Bim^-/-^* mice.

**Figure 3.**
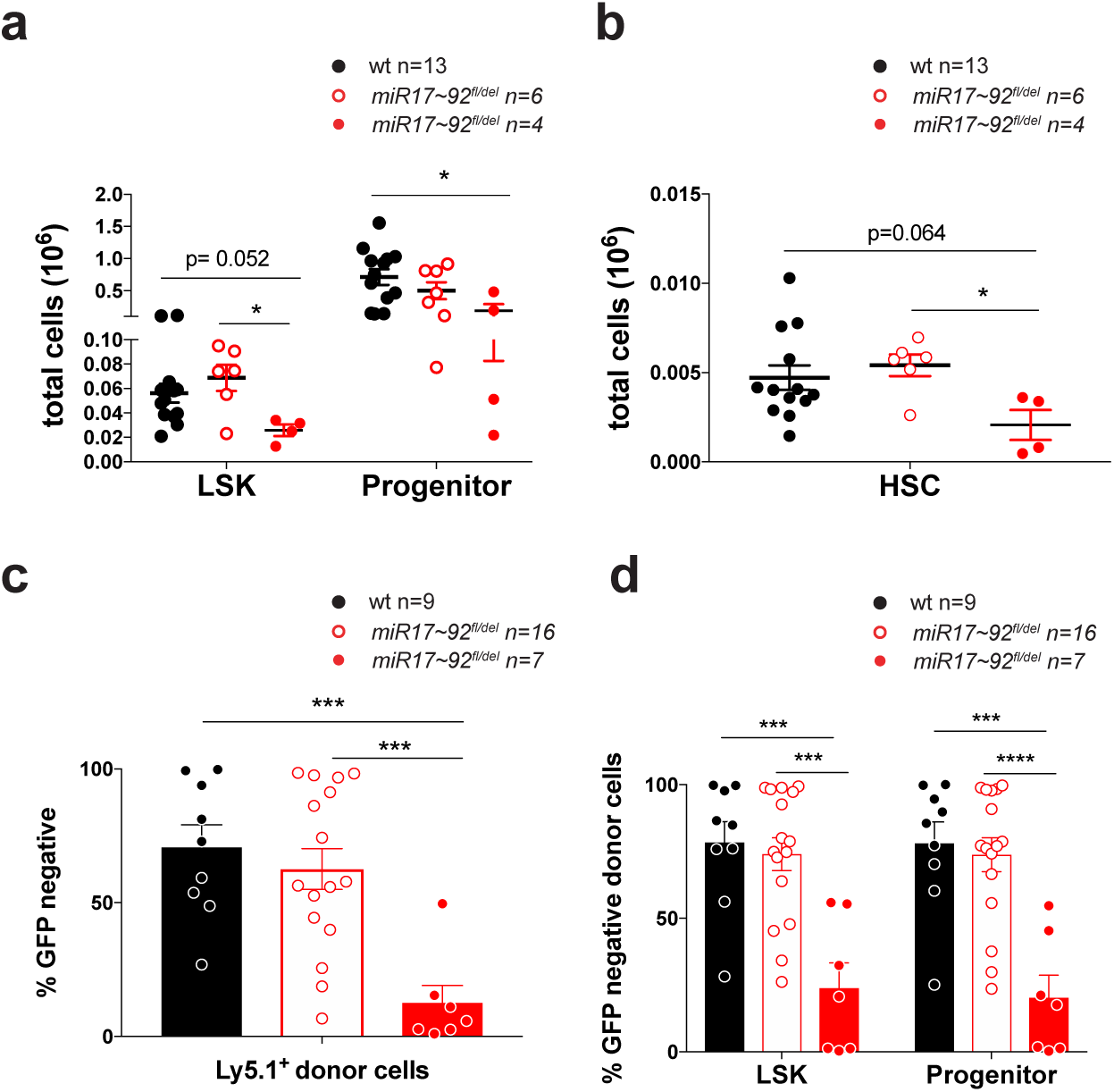
Induced deletion of the *miR17~92* cluster causes a loss of hematopoietic stem and progenitor cells. **(a)** LSK (Lineage-c-KIT^+^SCA-1^+^) stem cells and progenitor cells (Lineage-c-KIT-SCA-1^+^) were analyzed in E14 fetal liver cells from *miR17~92^-/-^*, *miR17~92^+/-^* and wt embryos by flow cytometry. **(b)** Hematopoietic stem cell (HSC) populations were identified in E14 fetal liver cells from *miR17~92^-/-^*, *miR17~92^+/-^* and wt embryos by staining for the SLAM markers. **(c-d)** C57BL/6 wt mice (Ly5.1+) were lethally irradiated and reconstituted with a 1:1 mixture of E14 fetal liver cells from UBC-GFP embryos (competitor cells) and E14 fetal liver cells from *miR17~92^-/-^*, *miR17~92^+/-^* or wt embryos (test cells). After 8-10 weeks, the fraction of reconstituting test cells was determined by gating on Ly5.1^+^GFP- cells (donor test cells) in FACS analysis. The percentages of GFP-negative test cells were determined for **(c)** donor cells and **(d)** LSK as well as progenitor cells. Data represent mean +/-SEM. *p<0.05, **p<0.01, ***p<0.001 (Student t test comparing *miR17~92^-/-^*, *miR17~92^+/-^* and wt).

### *imiR17~92* is critical for the survival of hematopoietic stem cells by restraining BIM-induced apoptosis

The fact that the induced deletion of *miR17-92* causes a reduction in all hematopoietic lineages could either be the result of independent loss of different immature and mature cell types or could be a consequence of the loss of common HSPC populations with consequent reductions in their differentiated progeny. To examine the latter, we first analyzed the HSPC populations in *miR17~92*-deficient E14 fetal liver cells. Remarkably, significant decreases in the numbers of LSK, HSC and progenitor cells were evident in *miR17~92^-/-^* fetal livers when compared to fetal livers from wt littermates (Figure 3a and b).

Competitive reconstitution assays using as test cells *miR17~92^-/-^*, *miR17~92^+/-^* or wt E14 fetal liver cells and E14 fetal liver cells from GFP embryos as competitor cells were performed to substantiate these findings. A significant competitive disadvantage to reconstitute the host hematopoietic system was observed for donor E14 fetal liver cells from *miR17~92^-/-^* embryos (Figure 3c). Detailed analysis of the hematopoietic stem/progenitor cell compartment revealed almost no contribution of the GFP-negative *miR17~92^-/-^* cells to the LSK and hematopoietic progenitor cell compartments, while the control donor cells (wt, *miR17~92^fl/-^*^)^ routinely contributed at least 50% to these HSPC compartments (Figure 3d and Supplementary Figure 3).

Next, we examined whether the loss of HSPCs caused by the induced deletion of *miR17~92* might be due to loss of the repression of pro-apoptotic BIM. Of note, microarray analysis revealed a significant negative correlation between *miR17hg (miR17~92* host gene) and *BCL2L11 (Bim)* expression in long-term hematopoietic stem cells (LT-HSC), short-term hematopoietic stem cells (ST-HSC), multipotent progenitor cells (MPP) as well as in lineage committed progenitor cells, including the common lymphoid progenitors (CLP), common myeloid progenitors (CMP), granulocyte/macrophage progenitors (GMP) and bi-potential erythroid/myeloid progenitors (BEMP) (Figure 4a). This suggests that *miR17-92* may be critical in HSPCs by restraining the levels of pro-apoptotic of BIM. To test this hypothesis, we again performed competitive bone marrow reconstitution assays and treated reconstituted mice 8-10 weeks post transplantation with tamoxifen to delete *miR17~92* in the test cells (strategy depicted in Supplementary Figure 2). We found that two months after *miR17~92* deletion almost no Lineage-c-KIT^+^SCA-1^+^ (LSK) stem cells or early progenitors (Lineage-c-KIT^+^SCA-1-) with deleted *miR17~92* were present in this competitive setting (Figure 4b). A more detailed analysis of the HSPC populations, using either the stem cell markers of the SLAM series ^30^ (Figure 4c) or the FLK series markers ^31^ (Figure 4d), revealed that deletion of *miR17~92* did not substantially affect the fitness of the LT-HSCs. However, deletion of *miR17~92* greatly diminished the competitiveness of the ST-HSCs as well as MPPs in comparison to the GFP+ (wt) competitor cells in the mixed bone marrow reconstitution assays. Strikingly, *miR17~92^fl/fl^;RosaCreERT2^Ki/wt^;Bim^-/-^* donor cells were not outcompeted by the GFP-expressing cells (Figure 4b-d). In fact the *miR17~92^fl/fl^;RosaCreERT2^Ki/wt^;Bim^-/-^* donor-derived cells contributed significantly, more than 50%, to several of these compartments, most likely due to a reduction in programmed cell death afforded by the complete loss of BIM. These results demonstrate that loss of *miR17~92* causes a reduction in ST-HSCs due to excessive BIM-induced apoptosis.

**Figure 4.**
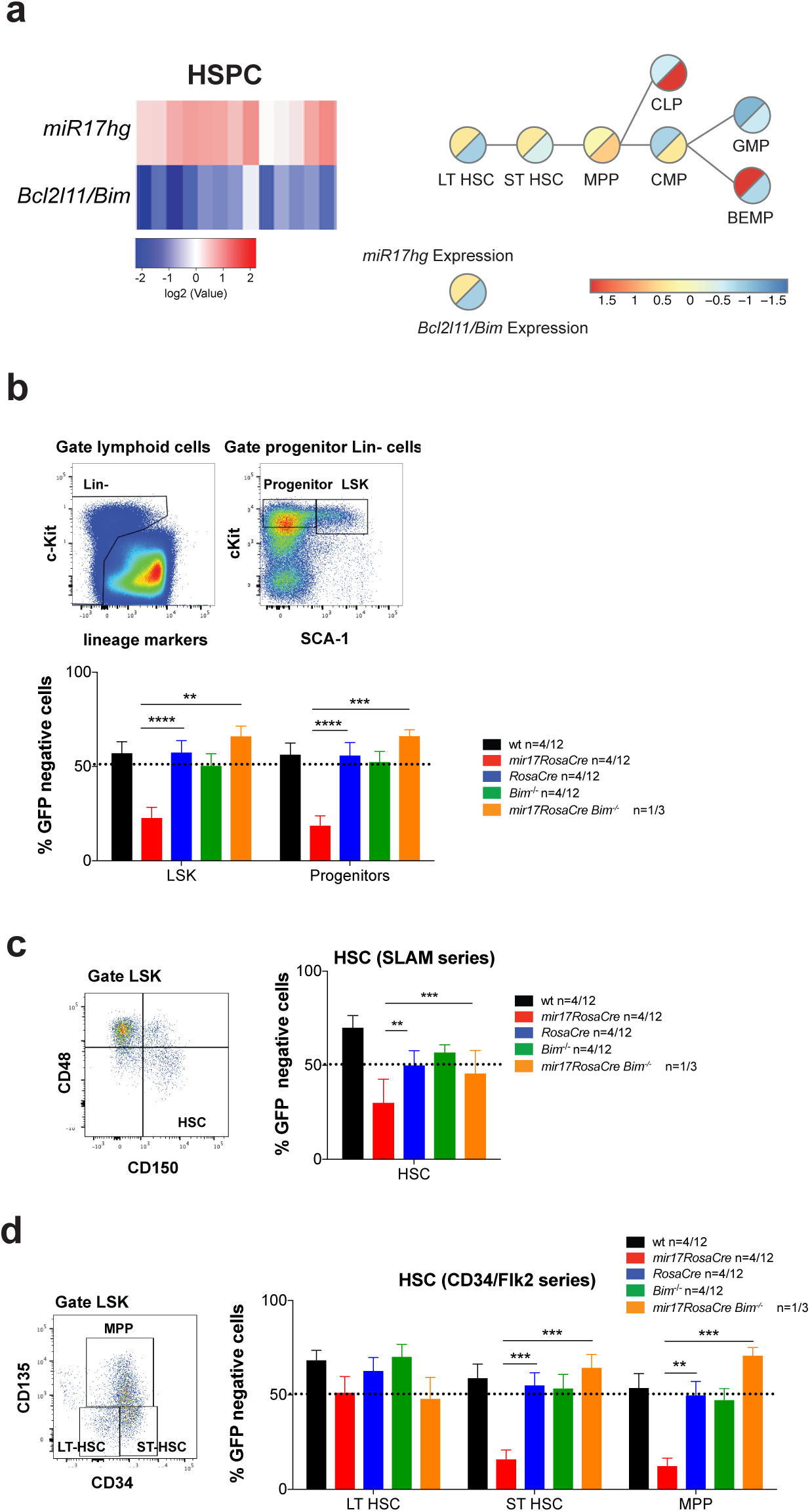
The depletion of hematopoietic stem and progenitor cells caused by the induced deletion of *miR17~92* is fully rescued by the loss of pro-apoptotic BIM. **(a)** Correlation analysis of *mir17hg (miR17~92*) and *BCL2L11 (Bim)* RNA expression in HSPC populations was determined by the examination of microarray data provided by immgen.org ^29^ (left panel). Schematic presentation of early hematopoietic stem/progenitor cell development with RNA expression analysis of *mir17hg* (*miR17~92)* and *BCL2L11 (BIM)* was performed as described (right panel) ^46^. **(b-d)** C57BL/6 wt mice (Ly5.1+) were lethally irradiated and reconstituted with a 1:1 mixture of bone marrow cells from UBC-GFP mice (competitor cells) and mice of the indicated genotypes (test cells). Reconstituted mice were treated with tamoxifen (3 doses oral gavage, 60 mg/kg/day) 8 weeks post-transplantation to activate the latent CreERT2 recombinase. After an additional 8-10 weeks, the hematopoietic stem/progenitor cell populations indicated were identified by staining with antibodies against cell type specific surface markers as indicated in the representative FACS blots. The percentages of GFP-negative cells (test cells) were determined for the indicated cell populations. Data represent mean +/- SEM. *p<0.05, **p<0.01, ***p<0.001, ****p>0.0001 (Student t test comparing *miR17~92^fl/fl^;RosaCreERT2^Ki/wt^* with *RosaCreERT2^Ki/wt^* and *miR17~92^fl/fl^;RosaCreERT2^Ki/wt^;Bim^-/-^* mice). Representative FACS blots for wt mice indicate the gating strategy and surface markers used. **(b)** LSK (Lineage-c-KIT^+^SCA-1^+^) cells and progenitor cells (Lineage-c-KIT-SCA-1^+^) were analyzed in the competitive bone marrow reconstitution assay. **(c)** Hematopoietic stem cells (HSC) were identified using cell surface staining for the SLAM markers in the competitive bone marrow reconstitution assay. **(d)** Long-term hematopoietic stem cells (LT-HSC), short-term hematopoietic stem cells as well multipotent progenitor cells (MPP) were analyzed using the FLK series markers in the competitive bone marrow reconstitution assay.

### *miR17~92* is critical for the survival of lineage committed hematopoietic progenitor cells by restraining BIM-induced apoptosis

The reduction in mature hematopoietic lineages and the HSC compartment caused by the deletion of *miR17~92* prompted us to also test the impact of induced deletion of *miR17~92* on lineage committed hematopoietic progenitor cells. This was again investigated by competitive bone marrow transplantation experiments and treatment of reconstituted hosts after 8-10 weeks with tamoxifen to inducible delete *miR17~92* (strategy depicted in Supplementary Figure 2). This analysis revealed that lineage committed pluripotent progenitors, including CLPs, CMPs, MEPs and GMPs, were substantially depleted upon the induced loss of *miR17~92* (Figure 5a). Consistent with this observation, we also observed reduced numbers of CLPs, CMPs, MEPs and GMPs in fetal livers cells from E14 *miR17~92^-/-^* embryos (Supplementary Figure S4a) and in the bone marrow of mice competitively reconstituted with a 1:1 mixture of fetal liver test cells from E14 *miR17~92^-/-^;RosaCreERT2^ki/wt^* embryos and competitor fetal liver cells from E14 GFP embryos (Supplementary Figure S4b). In all of these tests, early lineage committed progenitors of the myeloid, erythroid and megakaryocyte lineage were also significantly disadvantaged upon deletion of *miR17~92*. These included myeloid progenitors (pre-GMs and GMs) (Figure 5b), the erythroid colony forming units (pre-CFU-E, CFU-E) and the megakaryocyte/erythroid progenitors (MegE,) as well as the megakaryocyte progenitors (MK) (Figure 5c). Strikingly, BIM deficiency completely prevented the loss of all these progenitor populations caused by the inducible deletion of *miR17~92* (Figure 5). These results demonstrate that loss of *miR17~92* causes a reduction in lineage committed hematopoietic progenitor cells due to excessive BIM-induced apoptosis.

**Figure 5.**
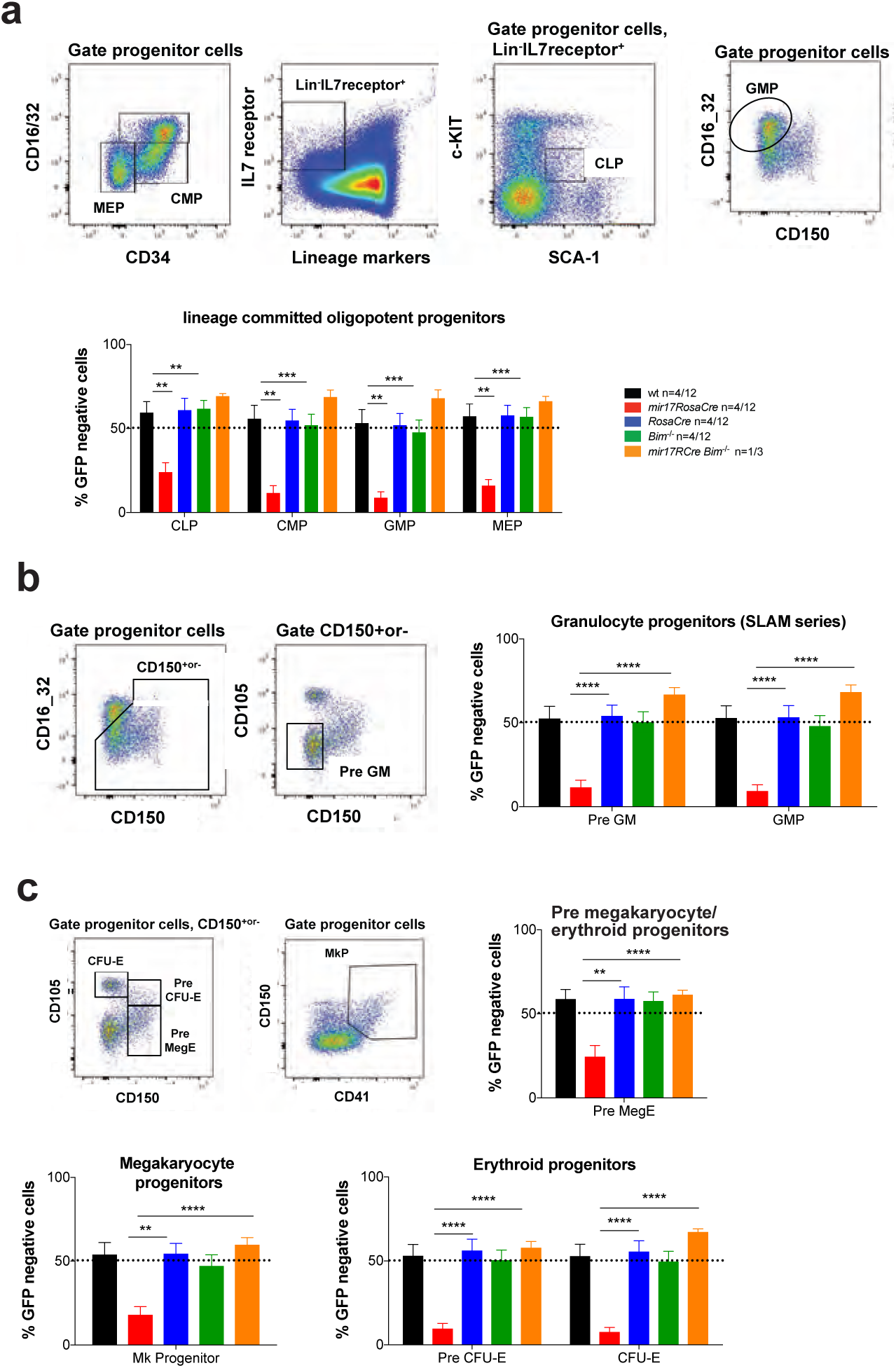
Deletion of *miR17~92* causes a substantial reduction in lineage committed hematopoietic progenitor cells and this can be completely rescued by concomitant loss of BIM. C57BL/6 wt mice (Ly5.1+) were lethally irradiated and reconstituted with a 1:1 mixture of bone marrow cells from UBC-GFP mice (competitor cells) and bone marrow cells from mice of the indicated genotypes (test cells). Reconstituted mice were treated with tamoxifen 8 weeks post-reconstitution to activate the latent CreERT2 recombinase. After an additional 8-10 weeks, the hematopoietic progenitor cell populations were identified by staining with antibodies against cell type specific surface markers as indicated in the representative FACS blots. The percentages of GFP-negative cells (test cells) were determined for the indicated cell populations. Data represent mean +/-SEM. *p<0.05, **p<0.01, ***p<0.001, ****p>0.0001 (Student t test comparing *miR17~92^fl/fl^;RosaCreERT2^Ki/wt^* with *RosaCreERT2^Ki/wt^* and *miR17~92^fl/fl^;RosaCreERT2^Ki/wt^;Bim^-/-^* mice). Representative FACS blots indicating the gating strategy and cell surface markers used are provided for wt mice. **(a)** Lineage committed progenitor cells, including CLP (common lymphoid progenitor), CMP (common myeloid progenitor), GMP (granulocyte/macrophage progenitor) and MEP (megakaryocyte/erythroid progenitor) populations, were examined in the competitive hematopoietic reconstitution assay. **(b)** Granulocyte progenitors, including the pre-GM and GM cell populations, were examined using the indicated markers. **(c)** Pre-megakaryocyte/erythroid progenitors (Pre-MegE), megakaryocyte (MK progenitors) and the erythroid progenitors, pre-CFU-E (colony forming unit erythroid) and CFU-E, were identified using antibodies against the indicated surface markers.

## Discussion

The *miR-17-92* cluster has been shown to play a critical role in the survival of immature as well as mature B lymphoid cells ^15^. Our present study reveals a previously unknown pro-survival function of *miR-17-92* in hematopoietic stem and progenitor cells (HSPCs). Remarkably, in these stem/progenitor cells loss of the pro-apoptotic BIM rescued almost all the defects caused by the deletion of the *miR17-92* cluster. While the conditional deletion of *miR-17-92* in adult mice did not impact on the LT-HSCs, all later stages, such as ST-HSC, MPP and lineage committed progenitors, depend on *miR17~92*-mediated suppression of BIM induced apoptosis for their survival and further differentiation.

Hematopoiesis is tightly regulated by several micro RNAs whose expression is dynamically controlled during differentiation and lineage commitment ^32,33^. To date the *miR17~92* cluster has been shown to regulate B cell development at the transition from the pro-B to the pre-B cell stage ^13,33^. Similarly, the ablation of DROSHA or DICER, the enzymes critical for micro RNA processing and maturation, causes similar defects in B cell development *^34^.* This demonstrates that the *miR17~92* cluster is the most relevant of all micro RNA clusters in the regulation of B cell development. Interestingly, transgenic overexpression of *miR17~92* resulted in an expansion of not only B lymphoid cells but also multipotent hematopoietic progenitors ^35^.

Our competitive bone marrow reconstitution assays demonstrated that the deletion of *miR17~92* severely impacts ST-HSCs and all of their progeny whereas LT-HSCs were not affected. During the process of hematopoiesis LT-HSCs are thought to divide asymmetrically, whereby the mother cell remains in a LT-HSC state in the stem cell niche, whereas the daughter cell exits this niche and enters the next stage of differentiation (reviewed in ^36^). This exit from the stem cell niche is accompanied by complex “re-programming” events, resulting in changes in transcriptional profiles that facilitate the proliferation, survival and differentiation into the distinct blood cell lineages. One of the most critical factors that is highly induced at the ST-HSC is c-MYC, a potent driver of cell growth and proliferation ^37^. The increase in c-MYC is a pre-requisite for the release of HSCs from the stem cell niche ^38^. Of note, c-MYC does not only induce cell growth and proliferation, but it also increases the predisposition of cells to undergo apoptosis particularly when the levels of growth factors and nutrients are limiting. Of note, this apoptosis is due in part to direct up-regulation of the pro-apoptotic BH3-only protein BIM by c-MYC ^39-41^. Moreover, it has been shown that anoikis – apoptosis induced by detachment of cells from their substrate (e.g. removal from the stromal cell niche) – is driven by JNK-mediated up-regulation of BIM ^42^. Hence, it can be argued that when HSCs leave the stem cell niche JNK cooperates with c-MYC to increase the levels of BIM. So, how is the pro-apoptotic activity of BIM opposed in the ST-HSC to safeguard their survival? One possible explanation is the concomitant transcriptional induction of the *miR17-92* cluster by c-MYC ^8,43^. Interestingly, BIM has been proposed to be one of the prime targets for the *miR17-92* cluster ^10,13^, based on the finding that loss of *miR17~92* causes an increase in the levels of BIM. This process might be involved in triggering apoptosis of stem cells that have exited the stem cell niche in the bone marrow.

Overexpression of *miR17~92* in B lymphoid cells can cause progressive lymphadenopathy, antibody mediated autoimmune disease and lymphoma ^44^. The authors of this study hypothesized that these abnormalities are driven exclusively by *miR17~92*-mediated repression of *Pten* as they only found a small reduction of BIM protein. However, this study only revealed a correlation and no functional (genetic) tests had been conducted, for example to show that loss of PTEN could phenocopy *miR17~92* over-expression. Of note, this study is not necessarily conflicting with our findings that loss of BIM achieves a complete rescue of the reduction in HSPCs that is caused by the induced deletion of *miR17~92* (Figure 3). Interestingly, earlier studies using *miR17~92^tg/tg^* and *miR17~92^-/-^* mice showed an inverse correlation between *Bim* and *miR17~92* expression in lymphocytes ^10,13^. Using the CD19-CRE or MB1-CRE conditional systems of transgene expression did not allow for the analysis of effects of *miR17~92* in HSPCs as they are only active from the pro-B cell stage of B lymphopoiesis, but not in HSPCs.

Taking all of the published data and our findings into account, we postulate that *Bim* induced apoptosis is the critical process that is restrained by *miR17~92* in HSPCs, including ST-HSCs, MPPs and lineage committed progenitors, with the loss of these progenitor populations caused by the induced deletion of *miR17~92* also affecting all of their differentiated progeny, such as B and T cells. The role of BIM in the defects caused by the loss of *miR17~92* in differentiated hematopoietic cells directly may be more complex. The extent of the rescue from cell death induced by *miR17-92* deletion that was afforded by the loss of BIM varied between different cell subtypes. For example, loss of BIM was able to completely rescue pro-B cells, pre-B cells and CD4-8- (DN) T cell progenitors in the thymus, whereas there was less pronounced rescue in granulocytes or macrophages. This is in line with the observation that BIM plays more prominent roles in the programmed death of lymphoid cells than myeloid cells ^45^. Remarkably, complete rescue from the impact of loss of *miR17~92* was observed in all hematopoietic stem and progenitor cells.

In conclusion, our studies reveal that *miR17~92* plays a critical role in the survival of HSCs and committed progenitors by restraining the expression of pro-apoptotic BIM. These findings will have implications for therapeutic strategies designed to target *miR17~92*. Moreover, it will be interesting to investigate whether *miR17~92* and BIM also play roles in the control of stem cell populations in other tissues, such as the colon or breast.

## Acknowledgements

We thank C Stivala, S Russo, J Mansheim, T Baldinger, T Kitson, C Gatt K McKenzie and G Siciliano for expert animal care; B Helbert and K Mackwell for genotyping; J Corbin and J McManus for automated blood analysis; E Tsui, V Orlando, K Weston, Y Hoang, C Tsui, S Ter for help with histology, S Monard and his team for help with flow cytometry and P Bouillet for providing *Bim^-/−^* mice. This work was supported by grants and fellowships from the Deutsche Krebshilfe (Dr. Milded-Scheel-post-doctoral fellowship to KB) the Australian National Health and Medical Research Council (NHMRC) (Project Grant 1145728 to MJH 1143105 to MJH and AS; Program Grant 1016701 to AS and Fellowship 1020363 to AS, Fellowship GNT1035229 to CAdG), the Leukemia and Lymphoma Society of America (LLS SCOR 7001-13 to AS and MJH), the Cancer Council of Victoria (1052309 to AS and Venture Grant MJH and AS), NHMRC Fellowship (1058344), NHMRC Program Grant (111357, (all to WSA) as well as by operational infrastructure grants through the Australian Government Independent Research Institute Infrastructure Support Scheme (361646 and 9000220) and the Victorian State Government Operational Infrastructure Support Program.

## Author contributions

KB, MJH and AS designed and conceived the study and experiments and prepared the manuscript. KB conducted and analysed the experiments. CAdG helped with the analysis of the microarray data. CDH and WSA provided advice for the analysis of the LT-HSC compartment helped with the experiments and data analysis and provided the antibodies and staining solutions.

## SUPPLEMENTAL MATERIALS

**Supplementary Figure S1**

**(a, b)** Mice of the indicated genotypes were treated with tamoxifen to activate the latent CreERT2 recombinase and thereby cause recombination of the floxed *miR17~92* alleles. Mandible bleeds were taken for hemograms at the indicated time points. The numbers of **(a)** monocytes, eosinophils, neutrophils, **(b)** hematocrit (HCT) and hemoglobin content (HGB) were determined using an ADVIA machine. **(c)** Immature and mature hematopoietic cell subsets from the bone marrow, spleen and blood were analyzed by flow cytometry at 1 or 6 months after tamoxifen induced CreERT2 mediated deletion of *miR17~92*. Data represent +/-SEM. *p<0.05, **p<0.01, ***p<0.001 (Students t test comparing *miR17~92^f/lfl^;Rosa26CreERT2^Ki/wt^* and *Rosa26CreERT2^Ki/wt^* or wt mice). **(d)** Histological analysis of H&E-stained sections of the spleen (right panel) and bone marrow (sternum, left panel) of wt, *Rosa26CreERT2^Ki/wt^* and *miR17~92^f/^lfl;Rosa26CreERT2^Ki/wt^* mice 1 month (upper panel) or 6 months (lower panel) after tamoxifen-induced CreERT2 mediated deletion of *miR17~92*. **(e)** PCR genotyping (detecting *miR17~92* wt, floxed and del bands) in bone marrow (left) and spleen cells (right) from wt (n=6), *miR17~92^fl/fl^* (n=5), *RosaCreERT2^Ki/wt^* (n=5) and *miR17~92^fl/fl^;RosaCreERT2^Ki/wt^* (n=10) mice at 6 months post treatment with tamoxifen to activate the latent CreERT2 recombinase.

**Supplementary Figure S2**

**Study design.** Lethally irradiated wt (C57BL/6-Ly5.1) mice were reconstituted with a 1:1 mixture of GFP-expressing wt (C57BL/6-Ly5.2) competitor bone marrow cells and test bone marrow cells with the genotype of interest, including *miR17~92^fl/fl^;RosaCreERT2^Ki/wt^*. Subsequent to the verification of hematopoietic reconstitution by FACS analysis of blood cells ~10 weeks post-transplantation, recipient mice were treated with tamoxifen to activate the latent CreERT2 recombinase and delete the *miR17~92* cluster in the test cells. After an additional 8-10 weeks, the hematopoietic progenitor cell populations were identified by staining with antibodies against cell subset specific surface markers. The percentages of GFP-negative cells (test cells) were determined for the indicated cell populations.

**Supplementary Figure S3**

C57BL/6 wt mice (Ly5.1+) were lethally irradiated and reconstituted with a 1:1 mixture of E14 fetal liver cells from UBC-GFP embryos (competitor cells) and E14 fetal liver cells from *miR17~92^-/-^*, *miR17~92^+/-^* or wt embryos (test cells). After 8-10 weeks, the fraction of reconstituting test cells was determined by gating on Ly5.1^+^GFP- cells (donor test cells) in FACS analysis. The percentages of GFP-negative test cells were determined for the indicated cell populations in **(a)** peripheral blood, **(b)** spleen, **(c)** bone marrow and **(d)** thymus. Data represent mean +/-SEM. *p<0.05, **p<0.01, ***p<0.001 (Student t test comparing *miR17~92^-/-^*, *miR17~92^+/-^* and wt).

**Supplementary Figure S4**

**(a)** Lineage committed progenitor cells, including CLP (common lymphoid progenitor), CMP (common myeloid progenitor), GMP (granulocyte/macrophage progenitor) and MEP (megakaryocyte/erythroid progenitor) populations, were examined in E14 fetal liver cells from *miR17~92^-/-^*, *miR17~92^+/-^* and wt embryos. **(b)** C57BL/6 wt mice (Ly5.1+) were lethally irradiated and reconstituted with a 1:1 mixture of E14 fetal liver cells from UBC-GFP embryos (competitor cells) and E14 fetal liver cells from *miR17~92^-/-^*, *miR17~92^+/-^* and wt embryos (test cells). After 8-10 weeks, the fraction of reconstituting test cells was determined by gating on Ly5.1^+^GFP- cells (donor test cells) in FACS analysis. The percentages of GFP-negative test cells were determined for the indicated cell populations. Data represent mean +/-SEM. *p<0.05, **p<0.01, ***p<0.001 (Student t test comparing *miR17~92^-/-^*, *miR17~92^+/-^* and wt).

## Methods

### Mice

Experiments with mice were approved by and conducted according to the guidelines of The Walter and Eliza Hall Institute Animal Ethics Committee. The generation of conditional *miR17~92^fl/fl^*, *RosaCreERT2^Ki/wt^*, *miR17~92^fl/fl^;RosaCreERT2^Ki/wt^* mice and the *Bim^-/-^* mice, all generated on a C57BL/6 or mixed C57BL/6x129Sv background, the latter backcrossed to C57BL/6 for >15 generations, has been described previously ^13,22,27^. The *miR17~92^fl/fl^;RosaCreERT2^Ki/wt^;Bim^-/-^* mice were generated by breeding *miR17~92^fl/fl^;RosaCreERT2^Ki/wt^* mice with *Bim^-/-^* mice. To activate the latent CreERT2 recombinase, mice were given 60 mg/kg tamoxifen (Sigma-Aldrich, Rowville, VIC, Australia) in peanut oil/10% ethanol each day for 3 days by oral gavage ^47^.

### Bone marrow reconstitution experiments

Bone marrow cells were harvested from both femora of mice at the age of 10 to 12 weeks, and single-cell suspensions were prepared. Bone marrow cells from such test mice of the genotypes of interest and competitor bone marrow cells from green fluorescent protein (GFP) transgenic mice (C57BL/6-Ly5.2 background), were mixed 1-to-1 in mouse tonicity adjusted saline (phosphate-buffered saline: PBS). From such cell mixtures, a total of 6 × 10^6^ cells per mouse were injected i.v. into 3 lethally irradiated (2 × 5.5 Gy, 3 h between doses) female congenic C57BL/6-Ly5.1 recipient mice 2 h after the second of g-irradiation. Eight weeks after transplantation, retro-orbital bleeds were taken to confirm successful hematopoietic reconstitution through determination of Ly5.2 positive cells by flow cytometric analysis.

### Blood analysis and flow cytometric analysis

Mandible bleeds were taken at the indicated time points and hemogram analysis was performed using the ADVIA system. Flow cytometric analysis of immature and mature hematopoietic cells in blood, spleen, bone marrow and thymus was performed as previously described ^48^. Hematopoeitic stem and progenitor cells (HSPC) were analyzed by staining with antibodies against specific cell surface markers: CD150-Bv421 (clone TC15-12F12.2, Biolegend), CD127-APC (clone A7R34, eBioscience), CD48-PECy7 (clone HM48-1, eBioscience), CD105-PE (clone MJ718, eBioscience), CD34-A647 (clone RAM34, BD), CD117-BV711 (clone 2B8, BD), CD16/32-PerCPCy5.5 (clone 2.4G2, BD), CD135-PE (eBioscience), CD41-PECy7 (clone MWReg30, BD), Sca1-A595, CD127-APCEF780 (eBioscience), CD9-A647, CD16_32-FITC, CD41-PECy7, CD2-A700, CD4-A700, CD8-A700, Gr1-A700, F4/8-A700, CD19-A700, B220-A700, Ly6G-A700, TER119-A700, Nk1.1-A700 (lineage markers). Specific HSPC populations were examined as described ^49^.

### Microarray analyses

Gene expression data from mature haematopoietic cells was taken from Immgen dataset (www.immgen.org) ^29^ and gene expression data for progenitor cells was taken from Haemopedia dataset (www.haemosphere.org) ^46^.

